# CONCENTRATION OF THIOUREA IS EFFECTIVE IN BREAKING THE DORMANCY OF POTATO VARIETIES

**DOI:** 10.1101/2020.12.02.407916

**Authors:** Sambat Ranabhat, Madhav Dhital, Ramkrishna Subedi, Ansu Adhikari, Binod Adhikari, Saroj Shrestha

**Affiliations:** Agriculture and Forestry University, Rampur, Nepal; Department of Horticulture, Agriculture and Forestry University, Rampur, Nepal; Plant Quarantine and Pesticide Management Center, Hariharbhawan, Kathmandu, Nepal

**Author notes:** Corresponding Author (SR). These authors contributed equally to this work (Madhav Dhital, Ramkrishna Subedi). These authors also contributed equally to this work (Binod Adhikari, Saroj Shrestha).

**Keywords:** Potato tubers (*Solanum tuberosum L*.), Dormancy breaking, Thiourea, Potato sprout

## Abstract

Potato germination is highly sensitive to ecological conditions. High altitude and low annual average temperature results in tuber dormancy and poor sprouting. Dormancy has become a significant constraint for lowering potato production that hinders the possibility of growing two crops per year. An experiment was conducted from February to April 2020, where two major potato varieties (Desiree and Cardinal) were treated with four concentrations of Thiourea (0, 1, 2, and 3%) in a two factorial completely randomized block design with three replications. Results showed that Thiourea has a significant effect on all observed attributes as per varieties of potato. For Desiree variety, Thiourea (1%) decreased breaking of dormancy by 22 days compared to control (Desiree*Thiourea 0%) and produced the longest average sprout of 7.36cm at 49 days after treatment (DAT). On the other hand, for the Cardinal variety, Thiourea (3%) decreased tuber dormancy by 27 days compared to control (Cardinal*Thiourea 0%) and produced a sprout of 7.75 cm at 49 DAT. In conclusion, for breaking dormancy and enhancing sprouting of potato varieties Desiree and Cardinal, Thiourea concentration of 1% and 3% is recommended, respectively.

**Author summary:** This work is the combined effort of all the authors; conceptualization and designing the plot experiments, S.R. and M.D.; performing the experiment and data collection, S.R. and A.A.; statistical analysis and preparation of presentation table and figure, S.R. and B.A.; writing the original draft and editing the whole manuscript, S.R., M.D., R.S., S.S., and A.A. All the author have read and agreed to the published version of the manuscript.

## Introduction

Potato *(Solanum tuberosum L)* is one of the vital food crops that is mostly grown in Nepal’s mid-hill region. In most developing countries such as Nepal, Potato is an important cash crop to resolve food insecurity and reduce poverty among smallholder farmers [1].

Several ecological conditions such as temperature, humidity, light, and soil play a key role during several developmental phases of potato such as germination, sprouting, vegetative growth, and maturity. In potato, the tuber germination and establishment stage is highly sensitive to severe cold leading to the tuber’s dormancy.

Dormancy is defined as the rest period under which sprouts fails to develop from any bud of tuber even though tuber is kept in ideal condition [2]. The tuber’s dormancy period is governed by several factors, i.e., potato cultivar, growing condition, storage duration, and tuber size [3,4,5,6]. The tuber’s dormancy results in low sprouting and poor establishment of the crop even in favourable growing conditions, which is one of the major causes of the decline in potato production. Also, more extended dormancy of tuber affects the possibility of cultivating potato more than once per year in various regions.

It is essential to break the dormancy of the potato tuber. Tuber treatment with a chemical such as Gibberellic acid, Thiourea, Ethylene Chlorohydrin, carbon disulfide, and Bromoethane can remove the tuber’s dormancy [7,8,9]. In recent years, Thiourea is used for breaking of dormancy of the potato tuber. Still, it is difficult to find research work that compares the effectiveness of various concentrations of Thiourea along with the potato varieties.

To address this knowledge gap, this research will help compare several levels of Thiourea concentration and determine effective one for breaking dormancy of potato tuber along with different varieties. Thiourea is defined as a catalyze inhibitor, which plays a crucial role in stimulating potato tuber germination. Application of Thiourea at appropriate concentration supports the germination process and develop multiple sprouts per eye of potato tuber [10,11]. Potato tuber producing a primary sprout length of at least 2mm is a reliable confirmation for breaking the dormancy period [12]. It was reported that potato tuber treated with Thiourea and hydrogen peroxide was found to have the early breaking of dormancy, i.e., 6 days and 10 days, respectively [13]. In Thiourea’s case, soaking the potato tuber in 1%, aqueous solution for one hour is suggested to break the tubers’ dormancy [7,10]. However, another report has shown that the treatment of freshly harvested mini tuber with 20 g/l Thiourea for 3 hours was effective than 30 g/l for dormancy reduction [14].

## Model

The research was carried out from 16^th^ February to 16^th^ April 2020. The experiment was based on two factorial CRD design with three replications. Two popular varieties of potato in Nepal (Desiree and Cardinal) were considered as 1^st^ factor. On the other hand, four Thiourea concentrations (0, 1, 2, & 3%) were taken as 2^nd^ factor.

## Methods

### Tuber collection

Freshly harvested potato tubers of two varieties were collected. Altogether 300 potato tubers of uniform shape and size were collected, i.e., 150 tubers of each type. The average weight of the tuber was 30-40 gm.

### Preparation of Thiourea solution

The Thiourea solution of different concentrations (1, 2, and 3%) was prepared by dissolving a calculated amount of Thiourea (Granules) in distilled water in separate plastic buckets. For a 1% solution, 30gm of Thiourea was dissolved in 3 litres of distilled water. Similarly, 60gm and 90gm Thiourea were dissolved in 3 liters distilled water for 2% and 3% solution.

### Details of treatment

The potato tubers of different varieties were treated with varying concentrations of Thiourea. Each treatment had ten potato tubers of uniform shape and size. The tubers were soaked in the solution for 2 hours. While for control, the tuber was soaked in distilled water for the same period. After that, the soaked tuber was air-dried until all the excess solution in the tuber surface was removed. Then, the tubers were placed in a plastic tray and kept in the darkroom.

### Bio-Metric observation

#### Days to the First emergence

The total days required to induce the emergence of the first sprout was taken as days to the first emergence. For this, potato tubers from each treatment combination were observed each morning to ensure the emergence of any sprouts. Only the sprout that reached a length of 2mm was counted as the first sprout from that particular tuber.

#### Breaking of Dormancy

Breaking of dormancy is considered by the number of days elapsing from the treatment until 80% of the tuber has at least one sprout equal to or longer than 2mm [12]. Here, a tuber from each treatment combination was inspected individually for producing the sprout of at least 2mm; once 80% of the tuber under observation had a sprout of at-least 2mm, then breaking of dormancy was calculated.

#### Sprouting length

The sprout’s length was measured using a scale equipped with millimeter reading for earlier data recording days. As the sprout length increased, a scale provided with centimeter reading was used. The average sprout length was measured regularly at an interval of 7 days up to 49 days of the experiment, starting from the first sprout initiation on the tuber.

#### Sprouting density

The total number of sprout per tuber was counted, and the average sprouting density was calculated. The tuber of each treatment combination was observed closely at an interval of 7 days. Those sprouts longer than 2mm from a particular tuber was counted, and sprouting density was determined.

### Data analysis technique

The data were collected at an interval of 7 days, and a total of 49 days of data was observed. The statistical analysis of data was done by using statistical packages, namely Microsoft Excel and R-studio. Duncan’s Multiple Range Test (DMRT) was used for mean separation and comparison at 5% significance level.

## Results and Discussion

### Early sprouting and breaking dormancy

Throughout the experiment, we can observe the significant effect of Thiourea’s different concentrations on dormancy breaking of tubers. The dormancy period was shorter in the tubers of both varieties treated with Thiourea compared to control. A Similar result was established by [10,13,15].

In tubers of Desiree variety treated with Thiourea (1%), sprouting started 6.33 days after treatment, whereas sprouting was started at 28 days for control. Moreover, the breaking of dormancy was achieved at 10.33 days after onset of treatment, which took 32.67 days for control tubers Table 1. On the other hand, tubers of Cardinal variety treated with Thiourea (3%) started sprouting at the 6.67^th^ day after the treatment while sprouting on control was initiated at 33.67^th^- day from the onset of treatment. In addition, breaking dormancy was found at 13.33^th^ day after the beginning of treatment, which was achieved after 40.67^th^ day for control Table 1. Furthermore, it was observed that Thiourea at any concentration shortens the dormancy period of tuber as compared to control

**Table 1:**
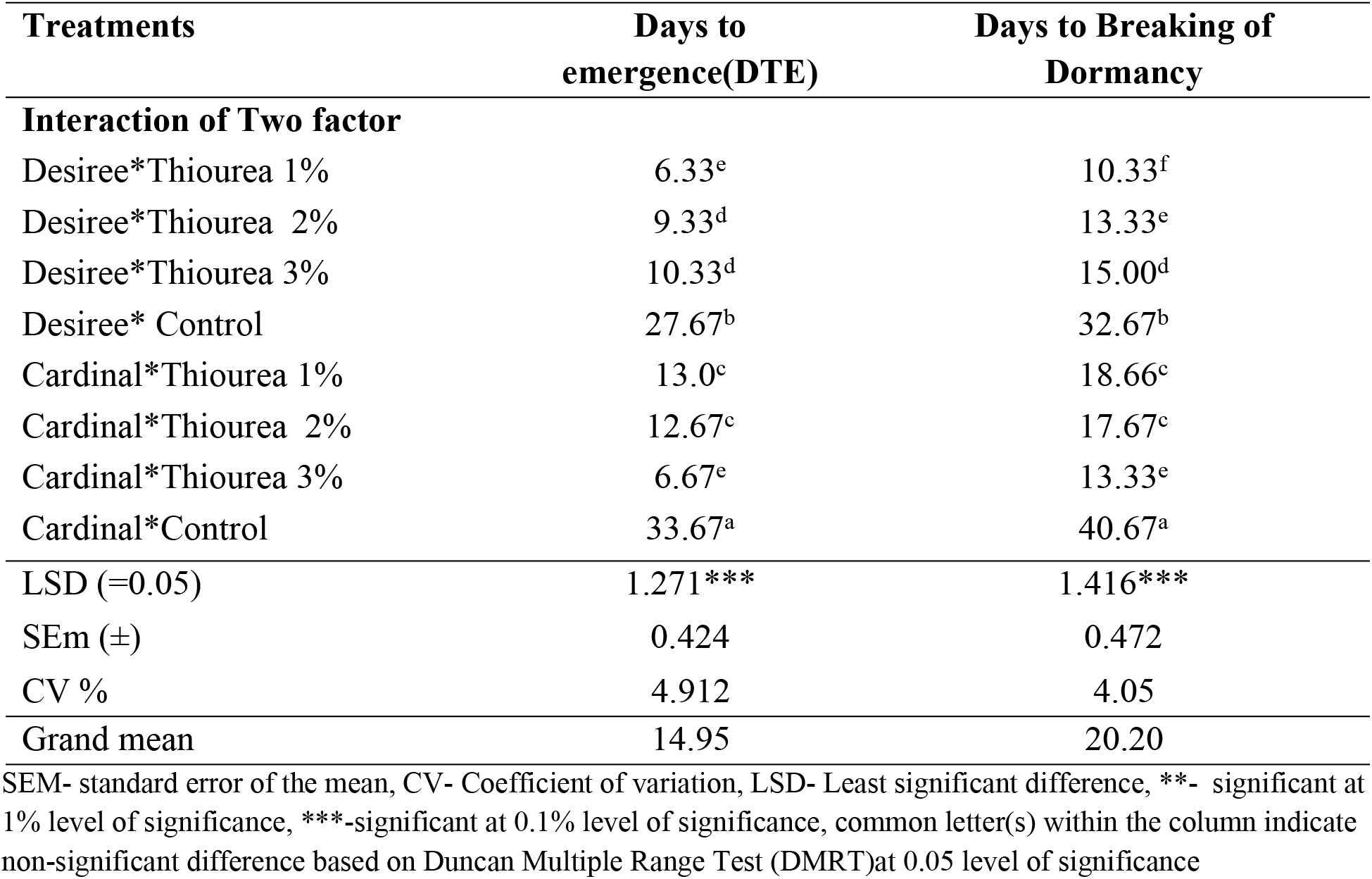
Days to emergence and breaking of dormancy as influenced by the interaction of variety and different concentration of Thiourea at Dailekh, Nepal (2020)

| Treatments | Days to emergence(DTE) | Days to Breaking of Dormancy |
| --- | --- | --- |
| **Interaction of Two factor** |  |  |
| Desiree*Thiourea 1% | 6.33<sup>e</sup> | 10.33<sup>f</sup> |
| Desiree*Thiourea 2% | 9.33<sup>d</sup> | 13.33<sup>e</sup> |
| Desiree*Thiourea 3% | 10.33<sup>d</sup> | 15.00<sup>d</sup> |
| Desiree* Control | 27.67<sup>b</sup> | 32.67<sup>b</sup> |
| Cardinal*Thiourea 1% | 13.0<sup>c</sup> | 18.66<sup>c</sup> |
| Cardinal*Thiourea 2% | 12.67<sup>c</sup> | 17.67<sup>c</sup> |
| Cardinal*Thiourea 3% | 6.67<sup>e</sup> | 13.33<sup>e</sup> |
| Cardinal*Control | 33.67<sup>a</sup> | 40.67<sup>a</sup> |
| LSD (=0.05) | 1.271*** | 1.416*** |
| SEm (±) | 0.424 | 0.472 |
| CV % | 4.912 | 4.05 |
| Grand mean | 14.95 | 20.20 |

SEM- standard error of the mean, CV- Coefficient of variation, LSD- Least significant difference, \*\*- significant at 1% level of significance, \*\*\*-significant at 0.1% level of significance, common letter(s) within the column indicate non-significant difference based on Duncan Multiple Range Test (DMRT)at 0.05 level of significance

In conclusion, for Desiree tubers, Thiourea (1%) shortens the dormancy period by 22 days compared to control. On the other hand, for Cardinal tubers, Thiourea (3%) shortens the dormancy period by 28 days compared to control Tables 1 and 2.

### Sprouting length

Treatment of the potato tubers with different Thiourea concentrations had a positive and significant increment in the average sprout length in both varieties. A similar result was conformed to those established by [14,15] on the increase in sprout length and thickness on tuber treatment with Thiourea.

For Desiree variety, using Thiourea with 1% concentration produced the longest sprout of cm length at 49 DAT which was significantly higher than other concentrations 0, 2, and 3%, which had sprout of 2.55, 6.02, and 5.72cm, respectively, at 49 DAT Fig 1. On the other hand, for Cardinal variety, Thiourea (1%) developed the longest sprout of 8.22 cm, which was statistically similar to 3% Thiourea, i.e., 7.75 cm at 49 DAT. Thiourea at a concentration of 2% and control produced shorter sprout of 6.58 cm and 1.92 cm at 49 DAT, respectively Fig 2.

**Fig 1:**
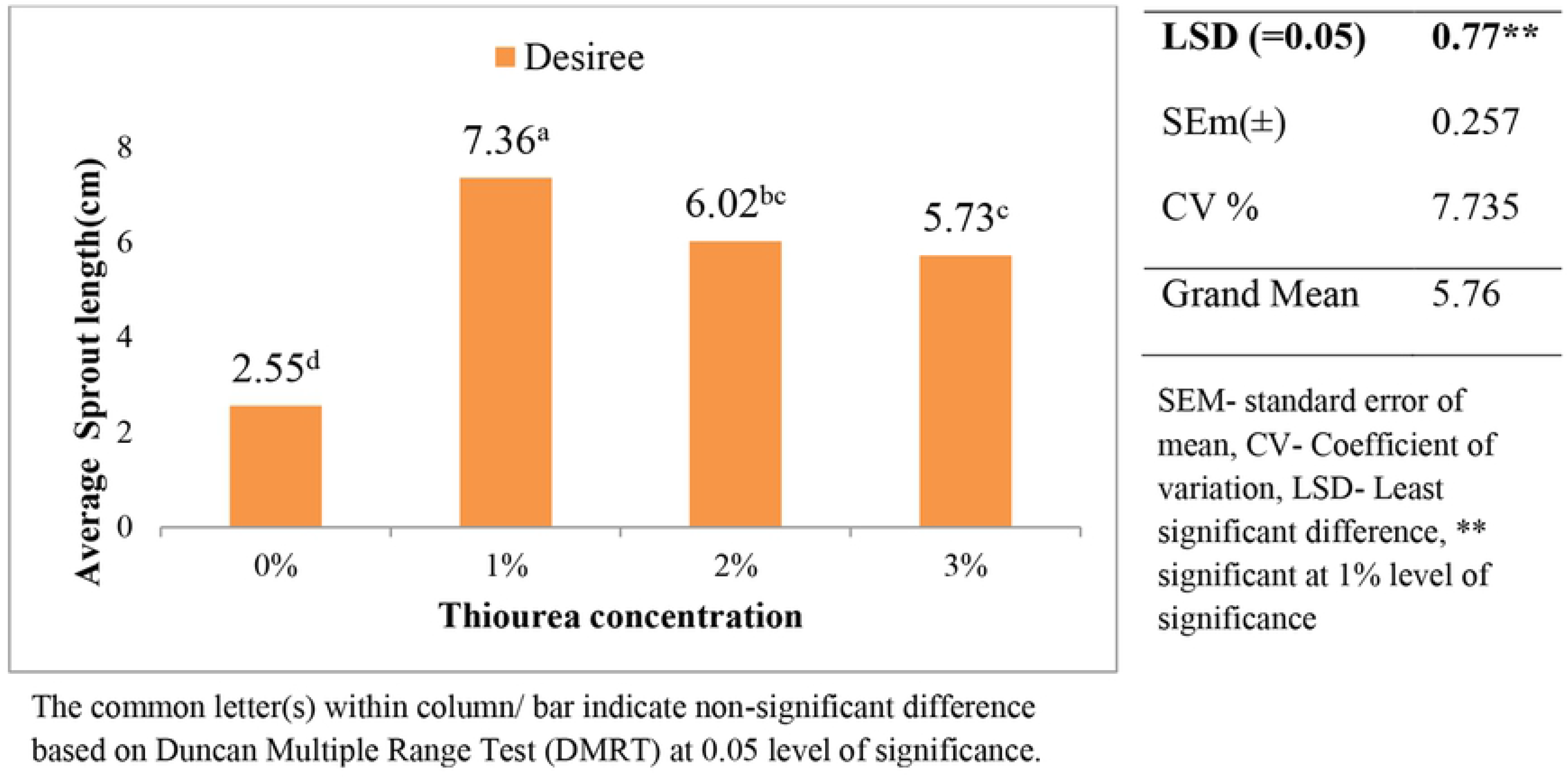
Effect of different Thiourea concentration on sprout length (cm) of Desiree potato at 49 days after treatment.

**Fig 2:**
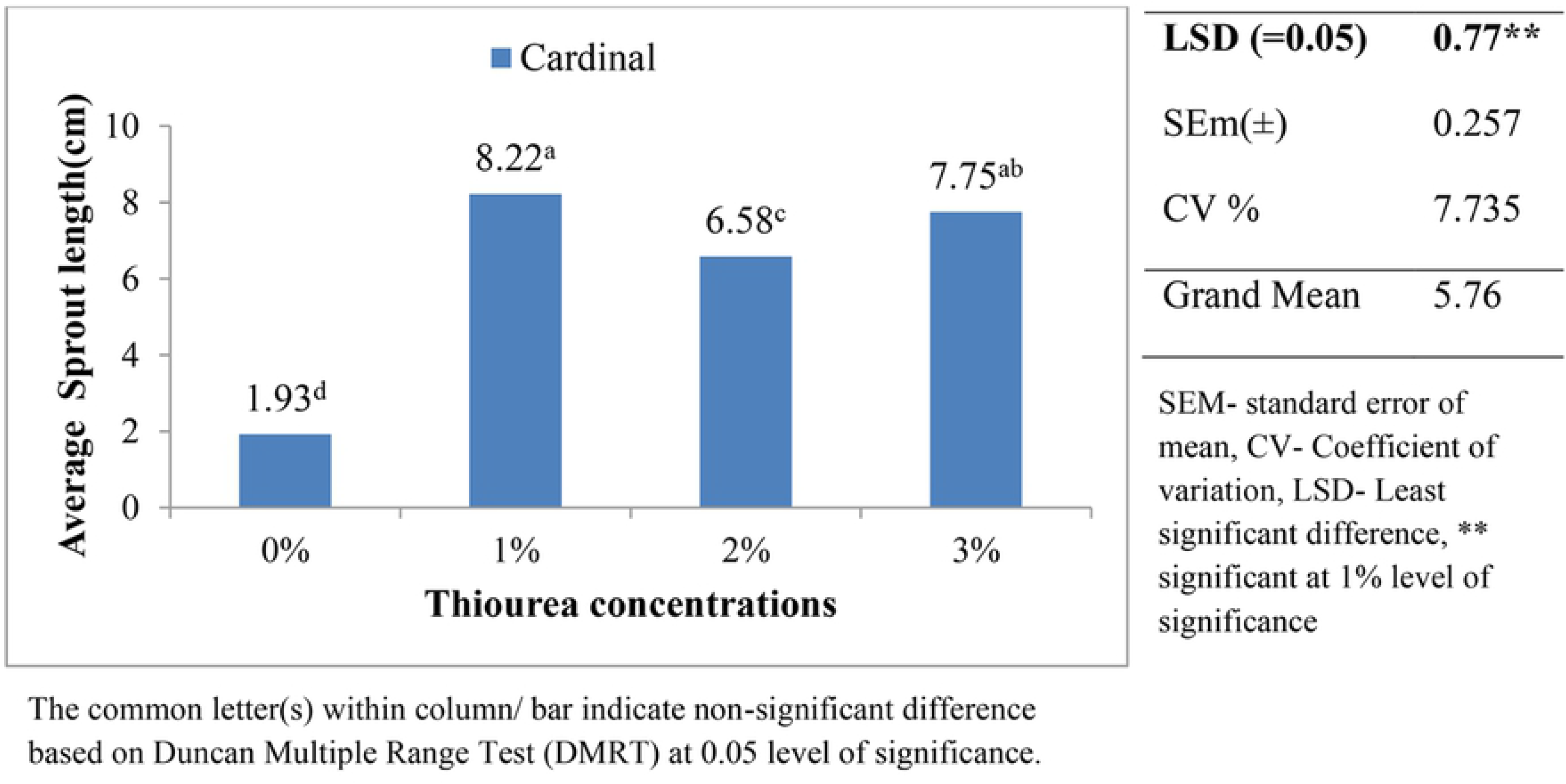
Effect of different Thiourea concentrations on sprout length (cm) of Cardinal potato at 49 days after treatment.

In conclusion, Thiourea (1% and 3%) was indistinctly superior to other Thiourea concentrations in Cardinal for developing the longest sprouts. While in the case of Desiree, Thiourea (1%) produced the longest sprout than the remaining concentrations, Fig 1 and 2.

### Sprouting density

Treatment of tubers with Thiourea of different concentrations showed a significant increment in sprout number per tuber for both varieties compared to control. The obtained result was similar to [11,14], who reported Thiourea treated tuber developed more sprout numbers than control.

Tubers of Desiree variety treated with Thiourea (1%) produced the greatest number of sprout, i.e., 4.13 sprouts/tuber, whereas control tubers made the least number of sprouts, i.e., 1.80 sprouts/tuber. Among all concentrations of Thiourea (1%) had significantly higher sprouting density for Desiree Fig 3. Whereas for tubers of Cardinal variety, Thiourea (3%) was found to produce significantly higher sprouting density, i.e., 1.91 sprouts/tuber compared to other concentration and lowest sprouts per tuber was developed by control, i.e., 1.13 sprouts/tuber Fig 4.

**Fig 3:**
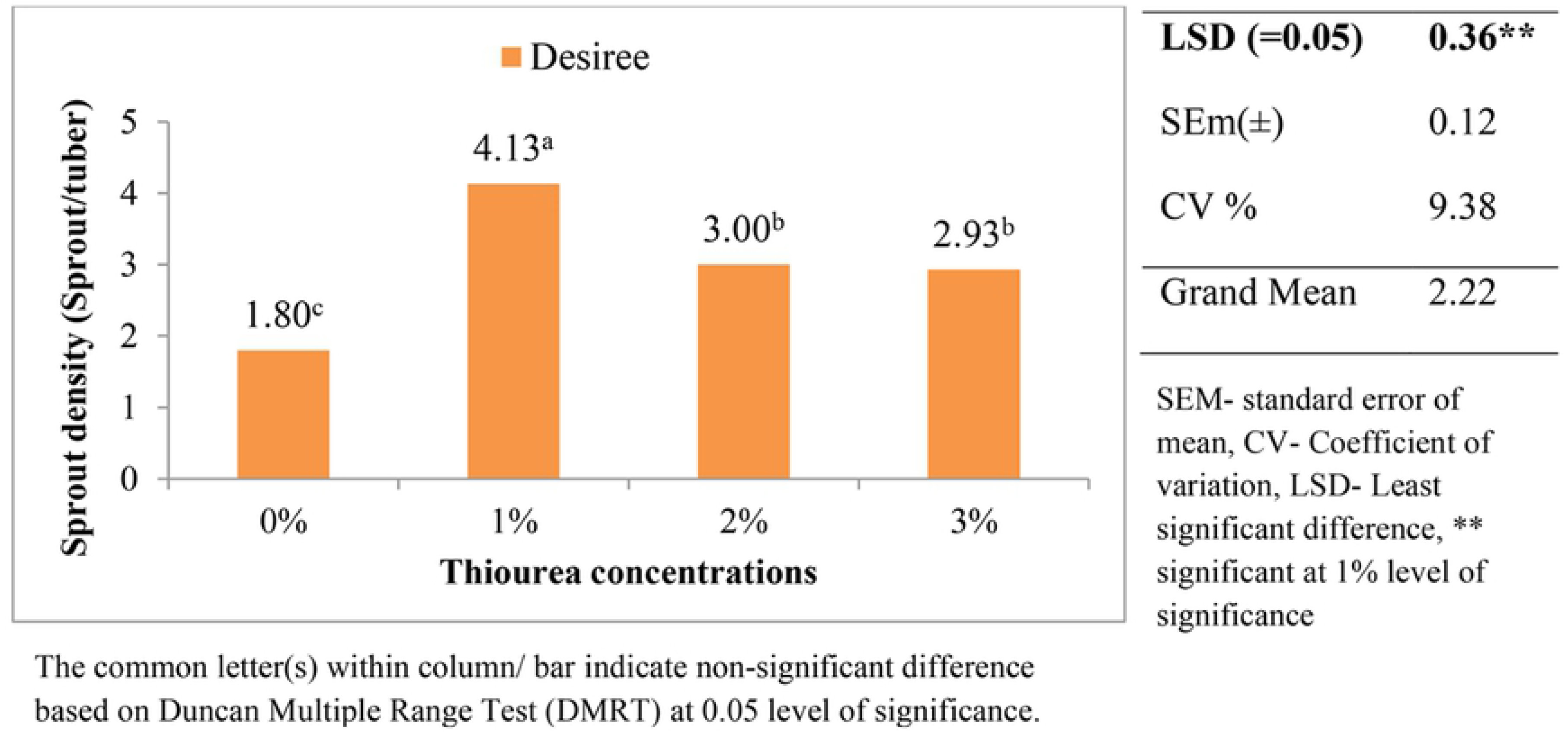
Effect of different Thiourea concentrations on sprout density (sprouts/tuber) of Desiree potato at 49 days after treatment.

**Fig 4:**
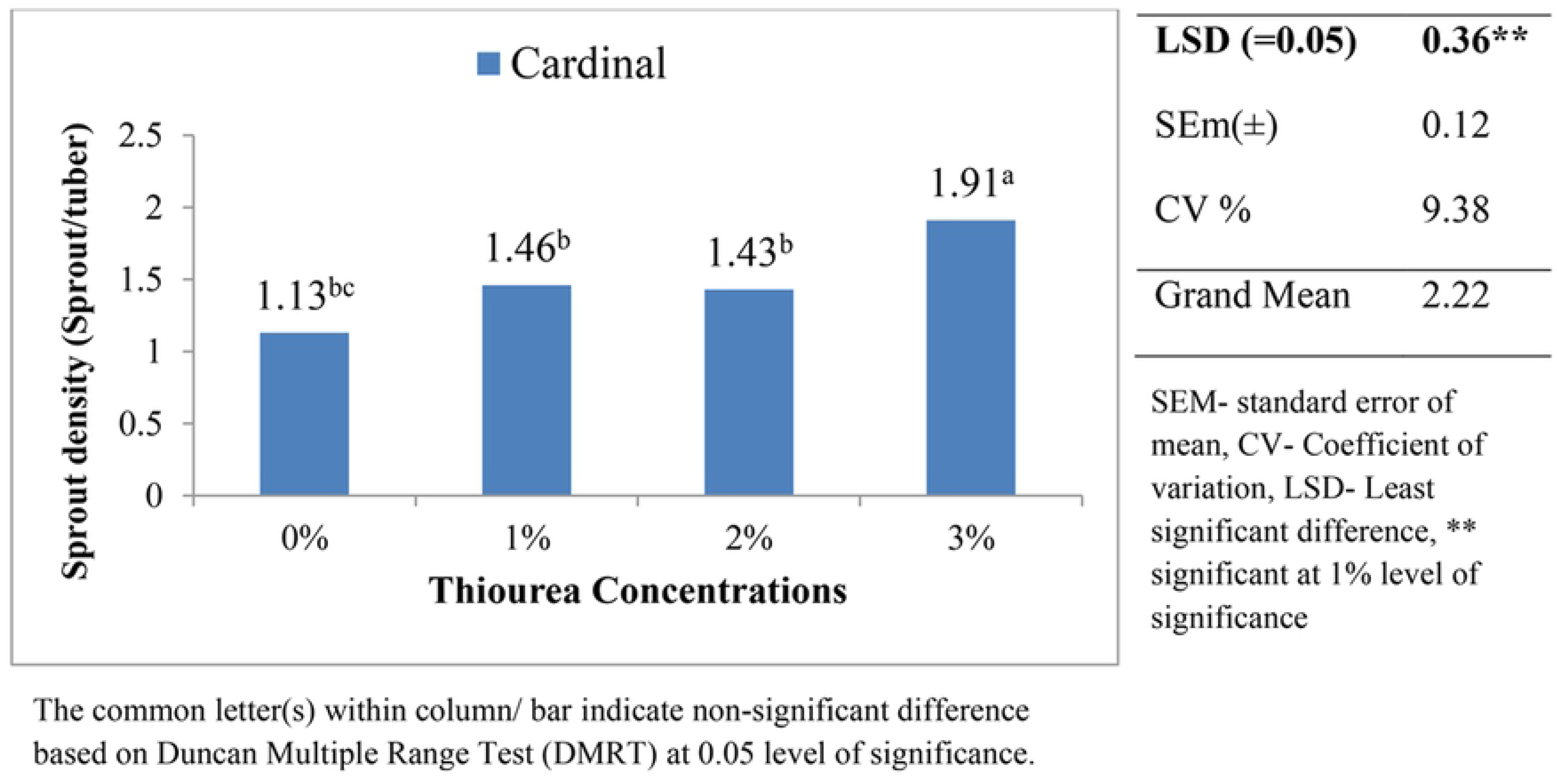
Effect of different Thiourea concentrations on sprout density (Sprouts/tuber) of Cardinal potato at 49 days after treatment.

In conclusion, Desiree and Cardinal Variety of potato, when treated with Thiourea of concentration 1% and 3% respectively, increased the sprouting number per tuber compared to control and other concentration Fig 3 and 4.

## Conclusion

In conclusion, among all treatment combination for Desiree variety, Thiourea (1%) was seen superior in a various parameter such as the emergence of first sprout (6.33^th^ day), breaking of dormancy (10.33^th^ day), sprout length (7.36cm) at 49 DAT and sprout density (4.13 sprouts/tuber) at 49 DAT. On the other hand, for cardinal variety, Thiourea (1%) was seen superior for a parameter such as a sprout length (8.22cm) at 49 DAT. However, the emergence of first sprout (6.67^th^ day), breaking of dormancy (13.33^th^ day), and sprout density (1.91 sprouts/tuber) at 49 DAT was found superior at Thiourea (3%).

Based on this study, it is recommended that a lower concentration of Thiourea, i.e., 1% was effective for dormancy breaking of potato tubers and enhancing sprouting behavior for Desiree variety. While for Cardinal variety, 3% Thiourea was found effective for dormancy breaking and improving sprouting behavior.

## Acknowledgement

I take great pleasure to express my sincere and intense sense of gratitude to Agriculture and Forestry University and the Prime Minister’s Agriculture Modernization Project (PMAMP).

